# Neural Network Models for Sequence-Based TCR and HLA Association Prediction

**DOI:** 10.1101/2023.05.25.542327

**Authors:** Si Liu, Philip Bradley, Wei Sun

**Affiliations:** Public Health Science Division, Fred Hutchinson Cancer Center, Seattle, USA; Herbold Computational Biology Program, Fred Hutchinson Cancer Center, Seattle, USA; Basic Sciences Division, Fred Hutchinson Cancer Center, Seattle, USA; Department of Biostatistics, University of Washington, Seattle, USA; Department of Biostatistics, University of North Carolina, Chapel Hill, USA

**Keywords:** T cell receptor, human leukocyte antigen, neural network, amino acid sequence, immune checkpoint blockade

## Abstract

T cells rely on their T cell receptors (TCRs) to recognize foreign antigens presented by human leukocyte antigen (HLA) proteins. TCRs contain a record of an individual’s past immune activities, and some TCRs are observed only in individuals with certain HLA alleles. As a result, characterising TCRs requires a thorough understanding of TCR-HLA associations. To this end, we propose a neural network method named Deep learning Prediction of TCR-HLA association (DePTH) to predict TCR-HLA associations based on their amino acid sequences. We show that the DePTH can be used to quantify the functional similarities of HLA alleles, and that these HLA similarities are associated with the survival outcomes of cancer patients who received immune checkpoint blockade treatment.

## Background

T cells play a critical role in the human immune response by recognizing antigens presented on the surface of cells by human leukocyte antigens (HLAs). This recognition is mediated by the T-cell receptor (TCR), a protein complex found on the surface of T cells. During an immune response, T cells that recognize a foreign antigen undergo clonal expansion, resulting in the production of a large number of T cells with identical TCRs. Once the target cells are cleared, the expanded clones contract to steady memory states and their TCRs can be detected years after the initial infection. A healthy person has a repertoire of tens of millions of TCRs [1]. Most of these TCRs are rare in the human population (or even private to an individual) because they are generated by a stochastic process. However, some TCRs are shared across individuals and they are known as public TCRs. These public TCRs likely arise because they are from the memory T cells responding to the same antigen (e.g., flu virus) in different individuals, or because they are generated with high probability [2]. Therefore, public TCRs can be used to infer immune response, such as infection history [1, 3].

HLA is the human version of major histocompatibility complex (MHC). The genetic locus encoding the HLA proteins is the most polymorphic region in the human genome [4, 5]. There are more than 15,000 classical HLA alleles [6]. We and others have shown that many public TCRs are restricted to certain HLA alleles by examining the co-occurrence of HLAs and TCRs in a subset of individuals [1, 7]. This approach is limited because it can only detect the associations of the HLAs and TCRs that are abundant in the population. It is desirable to have a more flexible method that can predict the association for any TCR-HLA pair. Such a method can be useful for many studies of immune response. For example, it can help define the functional similarities of HLA alleles and quantify the TCR-recognition capacity of a set of HLA alleles of a human being.

The challenge to train such a model is that both TCRs and HLAs are highly diverse. In addition to the diversity of TCRs and HLA alleles, each person can have up to 16 HLA alleles: up to six for three HLA class I (HLA-I) genes (two for each of HLA-A, HLA-B and HLA-C genes) and up to ten for three HLA class II (HLA-II) genes (two for HLA-DR, four for HLA-DP and four for HLA-DQ). To address this challenge, we propose a deep learning method named “DePTH”, which is short for **De**ep learning **P**rediction of **T**CR-**H**LA association. The main idea is to represent a TCR and an HLA by their amino acid sequences and learn their association by a neural network. This setup allows us to borrow information across HLAs and TCRs by exploiting their sequence similarity, and once the model is learnt, we can make a prediction for any TCR-HLA pair, as long as we know their sequences.

The TCR-HLA pairs for model training are built based on a set of 666 individuals[1, 7]. In later narrative, we refer to this data resource as “Emerson data”. We identified associated TCR-HLA pairs by their co-occurrence pattern. Our DePTH model learns to predict whether a TCR-HLA pair is associated or not based on their amino acid sequence information. We use all positive pairs and sample the negative pairs through a careful design to avoid potential bias.

Some recent works study the (TCR, HLA, peptide) three-way interactions by neural networks [8, 9, 10]. We focus on a different problem to model the associations between TCRs and HLAs, without being constrained to a specific peptide. To the best of our knowledge, the only other work that focuses on predicting TCR-HLA association is CLAIRE [11], which also uses a neural network model. Our work differs from CLAIRE in two aspects. First, DePTH represents an HLA allele by its sequence while CLAIRE treats HLA allele as a categorical variable, which limits its prediction to the HLA alleles included in the training data. Second, the training data are different. We identify associated TCR-HLA pairs by their co-occurrence [1, 7]. In contrast, CLAIRE uses associated TCR-HLA pairs from the database McPAS [12]. We show later that a model trained on one dataset does not generalize to the other, suggesting systematic difference of the two datasets. We argue that identifying positive TCR-HLA pairs by their co-occurrence is more objective. For example, it enables us to include more HLAs in the training process while the positive TCR-HLA pairs in McPAS are dominated by a few HLAs. Later in the Results Section, we show that when trained on co-occurrence data, DePTH outperforms CLAIRE, and when trained on McPAS data, DePTH makes more accurate prediction for all HLA-I alleles except one HLA allele that is the most abundant HLA allele in the McPAS dataset.

## Results

### An overview of DePTH

To collect training data, we use the co-occurrence of TCR-HLA pairs to select associated TCR-HLA pairs. Based on the first batch of 666 individuals from Emerson data [1], we treat the TCRs that appear in at least two individuals as public TCRs and assess the association between each public TCR and each HLA allele by onesided Fisher’s exact test. At p value cutoff corresponding to FDR 0.05 (Figure 1(A)), we selected 6,423 associated TCR-HLA pairs out of 742,832,595 TCR-HLA pairs involving HLA-I alleles and 11,037 associated TCR-HLA pairs out of 1,136,096,910 TCR-HLA pairs involving HLA-II alleles. These associated pairs are viewed as positive pairs.

**Figure 1.**
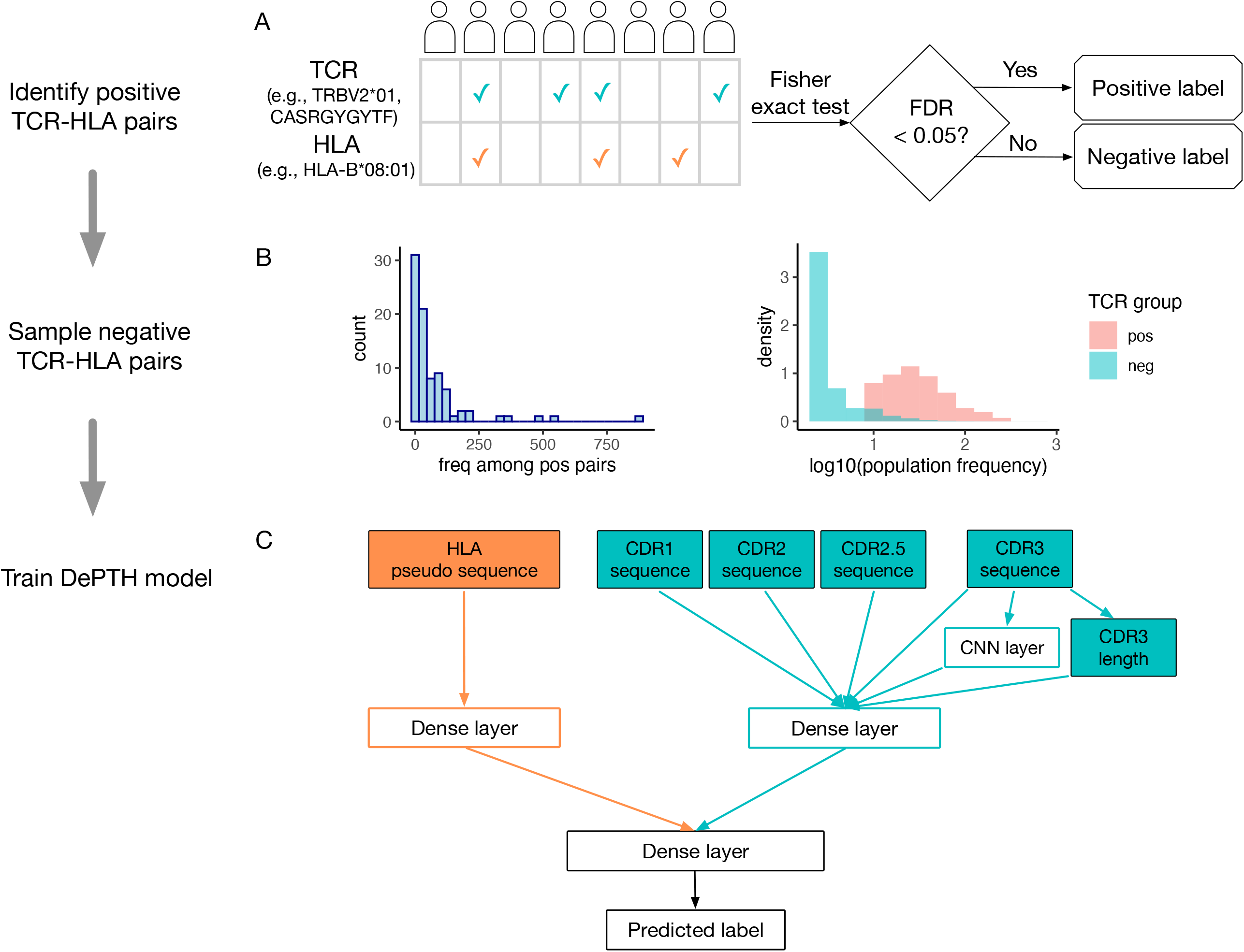
Summary for data generation and model training. (A) A toy example to illustrate the selection of positive TCR-HLA pairs by their co-occurrence. A check mark indicates the TCR or HLA is observed in an individual. The p value for the co-occurrence of a TCR-HLA pair is computed by one-sided Fisher’s exact test. To account for multiple testing across all TCR-HLA pairs, a TCR-HLA pair is considered associated if the corresponding False Discovery Rate (FDR) *<* 0.05. (B) The left panel is a histogram of the number of TCR-HLA pairs involving each HLA-I allele among the positive pairs identified from the Emerson data. The two overlaid histograms in the right panel show the population frequencies of TCRs that are associated HLA-B*01:01 or not, based on Emerson data. (C) Illustration of the components in the DePTH model. Each TCR is represented by its amino acid sequences from CDR1, CDR2, CDR2.5 and CDR3 parts, and each HLA is represented by its pseudo sequence (part of its sequence that interact with TCRs or antigens). DePTH learns to predict whether the pair is associated (positive) or not (negative). For HLA part, the encoded pseudo sequence is flattened and passed to a dense layer. For TCR part, the encoded CDR3 amino acid sequence is passed into a CNN layer. The output of the CNN layer is flattened and concatenated together with CDR3 length and the encoded and flattened amino acid sequences of CDR1, CDR2, CDR2.5, and CDR3. Then they are passed to a dense layer. The outputs of the two separate dense layers are concatenated together and passed to one or two dense layers before connecting to the final prediction.

When choosing the negative TCR-HLA pairs from those non-positive pairs, we follow two rules to avoid possible bias. The first rule is about the frequency of HLA alleles. Some HLA alleles appear much more frequently than other HLA alleles among the positive pairs (Figure 1(B), left panel). For example, HLA-B*08:01 appears in 878 positive TCR-HLA pairs and HLA-A*03:02 appears only in one. If we randomly sample negative pairs, the neural network may be biased to positive TCR-HLA pairs based on HLA allele frequency in training data. Therefore, we sample negative pairs so that the HLA frequencies in negative pairs are proportional to those among positive pairs. The second rule is to control the population frequency of TCRs. For each HLA allele, since the positive pairs are selected by Fisher’s exact test, the TCRs with higher population frequency are more likely selected (Figure 1(B), right panel). To avoid potential bias to score more prevalent TCRs as positives, for each HLA allele, we select the TCRs to form negative pairs so that their population frequencies are comparable to those of the TCRs in positive pairs. More details on training data preparation can be found in Sections 1.1 to 1.3 of the Supplementary Materials.

In our training data, positive and negative TCR-HLA pairs are encoded as 1 and 0, respectively. For each pair, the input for an HLA is its sequence at positions that contact the peptides or TCRs, and we refer to it as HLA pseudo sequence (see Section 2.1 of Supplementary Materials for details). A TCR is composed of an *α* chain and a *β* chain with more diversity on *β* chain. Most TCR data only include the *β* chain, which is the case for the Emerson data we use. Therefore we only consider *β* chain in this work, though our model can be easily extended to the situations where both *α* chain and *β* chain are available. The input for a TCR beta chain is its sequence in a few complementarity-determining regions (CDRs): CDR1, CDR2, CDR2.5 and CDR3, which are known to contact either the peptide or the HLA. CDR1, CDR2, and CDR2.5 are part of the V gene of a TCR (see Section 2.2 of the Supplementary Materials for details). CDR3, which is the most diverse region in a TCR, has variable lengths across TCRs. We pad all CDR3 sequences to length 27 by aligning them at both ends and padding the middle region by a character “.” [13] (Section 3.1 of the Supplementary Materials). The length of CDR3 before padding is included as an additional categorical input to the neural network.

Our neural network includes a dense layer to encode the HLA sequence, and a combination of a convolutional neural network (CNN) layer and a dense layer to encode the TCR sequence. CDR3 is the most diverse region in a TCR and it plays a more important role in TCR-HLA-peptide interaction, therefore we apply a CNN to CDR3 to capture non-position specific sequence motifs. The encoding of HLA and TCR are concatenated together and passed through one or two dense layers before final output is generated.

We split the data into training, validation and test sets. We train separate neural networks to predict TCR-HLA associations involving HLA-I alleles and HLA-II alleles, respectively. For hyperparameters, we consider amino acid encoding methods, number of dense layers and dropout probabilities in a dropout layer when predicting TCR-HLA association with concatenated inputs. We run cross-validation to choose the best hyperparameter setting. See Methods Section for details. Using the chosen hyperparameters, we train our model on training data, choose when to stop training based on the AUC (area under the ROC curve) on the validation data, save the model with the best validation AUC and evaluate its performance on test data.

### DePTH models achieve AUC around 0.8 on test TCR-HLA pairs

The best hyperparameter setting chosen from cross-validation for the situation of HLA-I alleles and that for the situation of HLA-II alleles are the same (see Supplementary Materials Table 1 for the hyperparameters chosen, and Supplementary Figure 1 for a graph showing the architecture of the model using the chosen hyper-parameters). The results are not sensitive to the hyperparameters. Most hyperparameters deliver similar and good performances.

Due to the inherent randomness of stochastic gradient descent algorithm, given the same set of hyperparameters, the predictions from different random seeds are not the same. We find that an ensemble prediction from multiple models (i.e., taking average prediction scores) delivers more robust and slightly more accurate predictions. We evaluate the ensemble prediction from *k* models where *k* increases from 1 to 100. The validation AUC became stable after ensemble size increased to around 20 (Supplementary Figure 2). In the following texts, all prediction scores and evaluation results from DePTH models are based on an ensemble of 20 models trained under 20 sets of random seeds, unless specified explicitly otherwise.

We evaluate the performance to predict TCR-HLA association by three metrics, AUC, sensitivity, and specificity. The sensitivity and specificity are computed based on a prediction score cutoff of 0.5 such that TCR-HLA pairs with prediction score greater than 0.5 are classified as associated pairs. DePTH delivers good classification accuracy, with AUC 0.82 and 0.79 for HLA-I and HLA-II alleles, respectively (see Figure 2(A) for the corresponding ROC curves). While the specificities are above 80% for both HLA-I and HLA-II alleles, the sensitivities are modest around 0.63. (Supplementary Materials Table 2).

**Figure 2.**
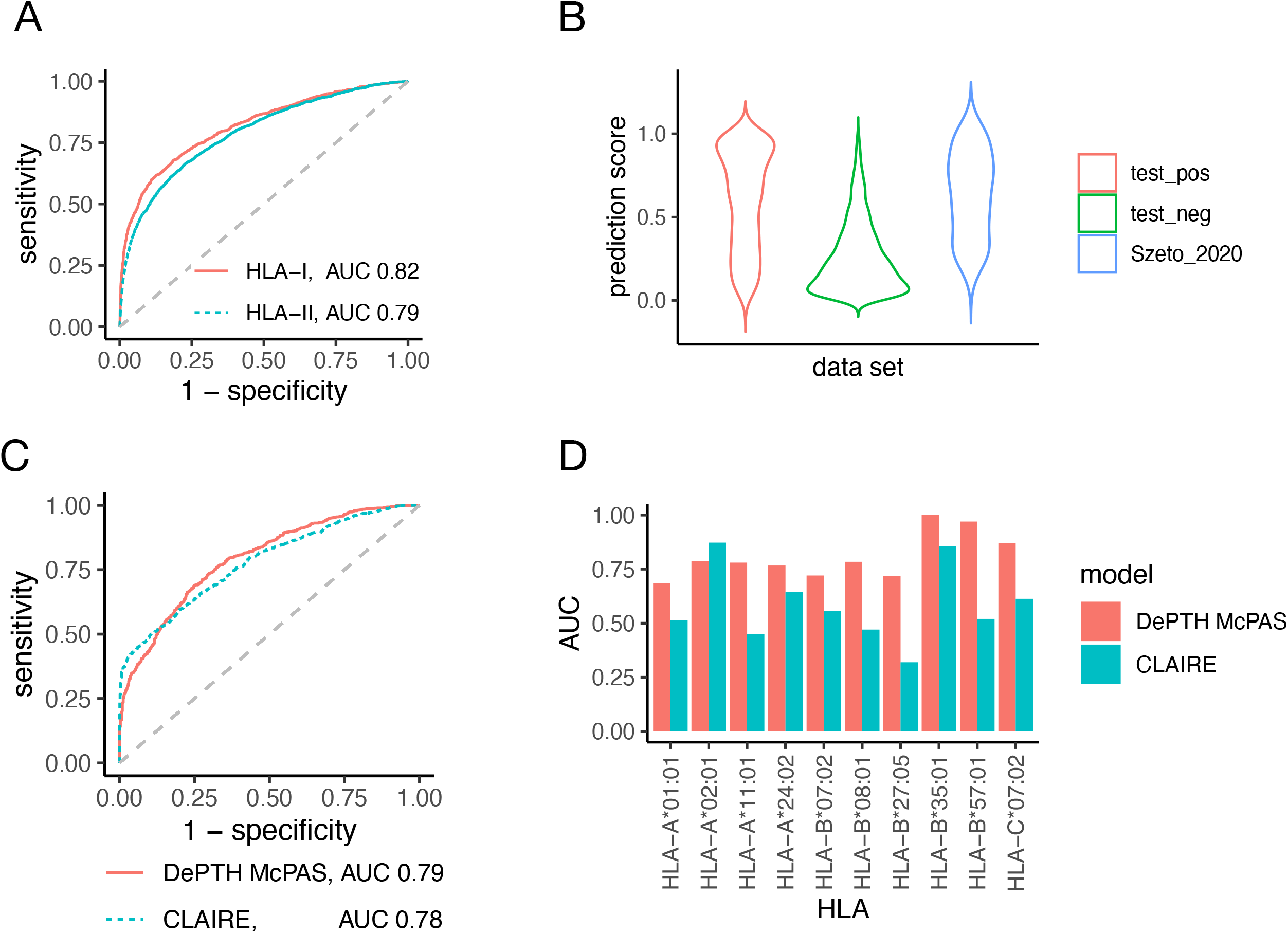
(A) and (B) demonstrate the performance of DePTH models trained on Emerson data. (A) the ROC curve of DePTH prediction on test data (part of Emerson data) for HLA-I and HLA-II, respectively. (B) Violin plots for the TCR-HLA association scores predicted by DePTH on three sets of TCR-HLA pairs: the positive pairs of Emerson test data, the negative pairs of Emerson test data, and the 54 external positive TCR-HLA pairs from solved TCR-pMHC-I structures provied by Szeto et al. [14]. The scores are given by DePTH model trained on Emerson training data. (C) and (D) compare the performance of DePTH and CLAIRE on McPAS test data while both models are trained on McPAS data. (C) ROC curves of prediction scores across all HLA-I alleles. (D) Comparison of allele-wise AUCs.

### DePTH can generalize to TCRs and HLAs unseen during training

In order to evaluate whether our model can generalize to TCR-HLA pairs that are not seen during model training and validation, we run three leave-one-out experiments. Each time, one of the top three most frequent HLA-I alleles that appear among the positive TCR-HLA pairs, namely HLA-B*08:01, HLA-B*07:02 and HLA-C*07:01, is left out. We chose these most frequent HLA-I alleles so that there are enough HLA-TCR pairs to obtain a reliable performance evaluation.

In each experiment, given the chosen HLA-I allele, for example, HLA-B*08:01, we construct the training, validation, and test data sets in the way such that (1) both positive and negative pairs in test data only involve HLA-B*08:01, while no pair in training and validation data involve HLA-B*08:01, and (2) there is no overlap between the TCRs from training or validation data with those from test data. To sample negative TCR-HLA pairs, we follow the same principles in terms of matching HLA frequency and TCR population frequency as those for the full data set. The ratio of the number of positive versus negative TCR-HLA pairs and the ratio of the sample size of training vs. validation pairs are the same as those listed in Methods Section. The hyperparameter setting is chosen from cross-validation. The test AUCs from these experiments range from 0.64 to 0.69 (Supplementary Materials Table 3), smaller than the AUC for the full data set, but they are still much larger than 0.5, which is expected by chance. Besides, the relatively lower performance is also likely due to reduced training sample size, since none of the training/validation pairs involves any HLA or TCR from test data.

### DePTH makes accurate prediction on TCR-HLA pairs from solved TCR-pMHC-I structures

We further evaluate DePTH by an independent data set provided by Szeto et al. [14], which consists of all the 81 TCR-pMHC-I complexes with solved structures deposited at Protein Data Bank by the year 2020. This data set provides a resource of TCR-HLA pairs that are known to bind together. We evaluate the performance of our model on this resource. After excluding the complexes involving non-human MHCs and additional processing (see Section 4 in the Supplementary Materials for more details), we end up with 54 unique TCR-HLA pairs. Based on prediction scores from our DePTH model trained on Emerson data, 67% of these TCR-HLA pairs are predicted to be positive at cutoff 0.5. When considering all cutoffs by an ROC, the AUC to distinguish these 54 TCR-HLA pairs from the negative pairs from Emerson test data is 0.87. We further examine the distributions of the prediction scores of these 54 TCR-HLA pairs, as well as the positive and negative pairs from Emerson test data (Figure 2B). These 54 TCR-HLA pairs have very similar prediction score distribution as the positive test set and higher prediction scores than the negative test set. This shows that DePTH model trained on Emerson data makes reasonable predictions for TCR-HLA pairs that do bind together.

### DePTH outperforms random forest

As a baseline model, in the situation of Emerson data involving HLA-I alleles, we train a random forest on the training data, where the inputs to random forest are the one-hot encoded amino acid sequences and CDR3 sequence length. The number of decision trees in random forest is increased in a step size of 100 until the validation AUC has not increased for two consecutive steps. The ensemble of decision trees giving the best validation AUC before the number of trees stops increasing is picked to give prediction on test data. Based on prediction scores given by random forest, the test AUC is 0.77 and the AUC to distinguish the 54 Szeto 2020 TCR-HLA pairs from negative pairs in Emerson test data is 0.68, both lower than corresponding results from DePTH. See Supplementary Figure 3 for the ROC curves of DePTH and random forest, and the violin plots for prediction scores.

### Comparison with CLAIRE

CLAIRE is a neural network model proposed by Glazer et al. [11] in 2022 for predicting the binding of TCR and HLA. It takes as inputs the HLA allele, V*β* gene and J*β* genes, and TCR *α* chain information as well when available. In this section, we discuss the major differences between CLAIRE and DePTH.

#### Difference in training data

We use the co-occurrence of HLAs and TCRs in a human population (the Emerson dataset [1]) to construct training data of associated TCR-HLA pairs. In contrast, CLAIRE mainly focuses on McPAS, which is a manually curated database [12] and relates TCRs with their associated MHCs based on published experimental data.

We show that the model trained on one dataset (either Emerson or McPAS dataset) does not generalize to the other. Specifically, we train DePTH models on one of the two datasets, and then make prediction on the other dataset. For McPAS data, the training, validation and test sets used for developing CLAIRE were obtained from the github repository for CLAIRE [11]. The process of training DePTH on the processed McPAS data is similar to that of training DePTH on Emerson data (See Sections 5.1-5.2 of the Supplementary Materials for more details).

Our model trained on processed McPAS data for HLA-I class achieves AUC 0.79 on McPAS test data, with sensitivity 0.72 and specificity 0.71. However, if we evaluate it on the Emerson test data, the AUC is only 0.52, which is close to random guess. On the other hand, if we evaluate the DePTH model trained on Emerson data by making predictions on the McPAS test data, the AUC is also 0.52. These results show that the model trained on one data source does not generalize to the other data source. This is consistent with the observation by Glazer et al. [11] that the CLAIRE model trained on McPAS data does not generalize to Emerson data. A possible reason is the TCR-HLA pairs from McPAS data might have higher binding affinity, as conjectured by Glazer et al. [11].

#### Difference in encoding the HLA and TCR input

DePTH takes the pseudo sequence of an HLA allele (i.e., amino acid sequence at the positions that interact with a peptide or a TCR) as input. This choice allows DePTH to be applied to any HLAs with sequence information. In contrast, CLAIRE treats HLA as a categorical variable, which limits its ability to generalize to unseen HLAs. Furthermore, for the V gene of a TCR, DePTH takes its amino acid sequence in CDR1, CDR2 and CDR2.5 as input features, which allows generalization to previously unseen V genes. In contrast, CLAIRE treats V gene as a categorical variable.

#### DePTH outperforms CLAIRE on most HLAs when trained on the same data

To show the prediction performance difference related to the encoding of input features, we compare the DePTH trained on McPAS training data (we refer to this model as DePTH McPAS) with CLAIRE model on McPAS test data. The prediction scores from CLAIRE model are obtained by submitting the test data files to the CLAIRE model provided by Glazer et al. [11] on a web server. In terms of overall AUC based on prediction scores on all test pairs, DePTH McPAS achieves 0.79, which is slightly better than 0.78 achieved by CLAIRE model, with ROC curves shown as in Figure 2(C). Furthermore, we focus on the 10 HLA-I alleles that each appears in at least one positive TCR-HLA pair and at least one negative pair in test data. For each HLA-I allele, we compute the allele-wise AUC based on prediction scores by DePTH McPAS or CLAIRE. DePTH McPAS has higher AUC than CLAIRE in 9 out of the 10 HLA-I alleles (Figure 2D), except for one, HLA-A*02:01, which is the most frequent HLA-I allele among the TCR-HLA pairs used for training CLAIRE and occupies 44% of TCR-HLA pairs. While for allele HLA-B*27:05, which is a much less frequent HLA-I allele and is observed in only ∼1% of these TCR-HLA pairs, AUCs from DePTH McPAS and CLAIRE are 0.72 and 0.32, respectively. For rare HLA alleles, DePTH has the advantage that it can learn from other HLA alleles with similar pseudo sequences.

The above results from DePTH McPAS are from an ensemble of 20 models trained with 20 sets of random seeds. Since ClAIRE model is a single model instead of an ensemble, to compare the two approaches on the level of single model, we also compare the results of one DePTH McPAS model trained with one set of random seeds with those of CLAIRE. The single DePTH McPAS model gives overall test AUC 0.76, which is lower than the 0.78 achieved by CLAIRE, with ROC curves shown in Supplementary Figure 4(A), but the trend of allele-wise AUC comparison (Supplementary Figure 4(B)) is consistent with the case of model ensemble, where the allele-wise AUC from the single DePTH McPAS model is higher than that from CLAIRE for all 9 HLA-I alleles, except for HLA-A*02:01.

### HLA similarity metrics defined based on DePTH prediction are associated with survival outcome of cancer patients treated with Immune Checkpoint Blockade (ICB)

DePTH allows us to transform an HLA from the space of amino acid sequences (input of DePTH) to the space of TCRs, since the output of DePTH gives the association between this HLA and a set of TCRs. We use such TCR-HLA associations to quantify HLA similarities. Since the outcome is cancer related, we compile a list of TCRs from CD8+ T cells with cancer reactive signatures, using a dataset with single cell TCR and gene expression data [15]. We first identify cancer-related CD8+ T cells using known gene expression signatures [16, 17, 18, 19], and then extract their TCRs. See Section 6.1 of the Supplementary Materials for more details. We define two types of metrics, one is between-individual HLA-I distance and the other one is individual-level HLA-I heterozygosity metric.

#### Between-individual HLA-I distance

Chowell et al. [20] reported a dataset containing both HLA-I information and the survival outcome for 1,535 advanced cancer patients treated with ICB. While we have identified pseudo sequences for all HLA alleles in Emerson data, some patients in Chowell dataset have HLA-I alleles that do not belong to the 85 HLA-I alleles from Emerson data. For each of these HLA-I alleles, we match it with the first allele in the group (at two-digit resolution) and use the pseudo sequence of the latter instead. See Supplementary Materials Section 6.2 for more details. We exclude patients with HLA-I alleles for which we cannot not find a match, and this reduces the total sample size from 1,535 to 1,443.

We apply DePTH to score the associations between each HLA-I allele from this population and the set of potentially cancer-related CD8+ TCRs (see Section 6.2 of the Supplementary Materials for details), and select the set of TCRs associated with each HLA-I allele by a cutoff 0.5. For each patient, we collect six sets of TCRs, one for each HLA-I allele of the patient (Figure 3(A)). If an HLA-gene is homozygous (e.g., two HLA-A alleles are the same), the two alleles are still considered as two entities in our computation. We define an individual-level TCR set by taking the union of these six TCR sets. Given two individual-level TCR sets, denoted by *A* and *B*, we compute a between-individual HLA-I distance as

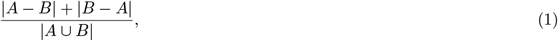

which is the number of TCRs belonging to only one of the two sets divided by the number of TCRs belonging to either set (Figure 3(B)), and is the same as 1 Jaccard similarity index between these two sets. We refer to this distance as “dist DePTH breadth” since it is based on prediction scores from DePTH model and the breadth of TCR set. Using this approach, we calculate a distance matrix for all individuals. Next, we assess the association between this distance matrix and survival outcome by “MiRKAT-S”, which is originally developed to assess the association between survival outcome and distance defined by microbiota [21].

**Figure 3.**
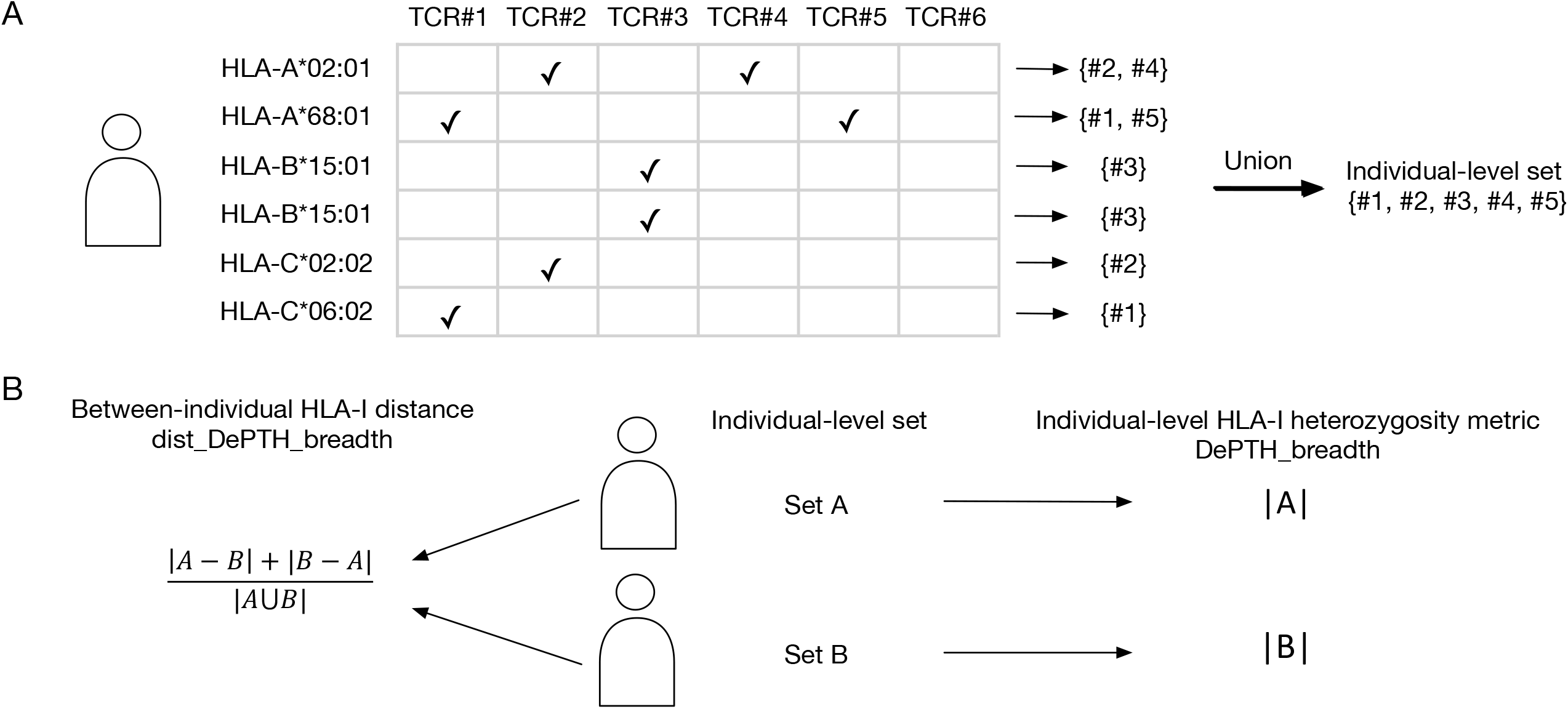
A toy example to illustrate the calculation of “dist DePTH breadth” and “DePTH breadth”. The top panel of the figure demonstrates the selection of the TCRs that are associated with at least one of this individual’s HLAs. In the HLA vs. TCR table, a check mark indicates predicted association between an HLA (each row) and a TCR (each column). The bottom panel of the figure illustrates the calculation of between-individual distance (“dist DePTH breadth”) and individual-level HLA-I heterogeneity metric (“DePTH breadth”). |*A*| and |*B*| are the sizes of the sets *A* and *B*. The between-individual distance is used for kernel regression and the individual-level metric is used for Cox regression.

We also define two other distances. One distance is “dist_DePTH_cor”, for which the between-allele distance is 1 - Spearman’s correlation of TCR-HLA association scores (predicted by DePTH) for two HLA alleles. The other distance is “dist_AA”, for which the between-allele distance is based on the values on Blosum62 matrix [22] corresponding to the amino acids on each aligned position of two HLA-I pseudo sequences (see more details Methods Section), and thus it is a metric purely based on amino acids. Since each individual has 6 HLA-I alleles, we need to transform distances between HLA-I alleles to distances between individuals. Towards this end, we apply optimal transport [23] to find optimal match of HLA-I alleles between two individuals and calculate between-individual distance. See Methods Section for more details. Using similar methods and CLAIRE prediction results, we also create two other between-individual distances “dist_CLAIRE_breadth” and “dist_CLAIRE_cor”.

For all five between-individual distances, we assessed their associations with survival outcome by MiRKAT-S, with and without covariates. As shown in Figure 4(A), at p-value cutoff 0.05, “dist_DePTH_breadth” shows significant association with survival outcome when no covariate is included, and the significance remains when age, log-transformed mutation burden, and drug type (CTLA-4, PD-1, or both) are included as covariates. “dist_DePTH_cor” gives p-value around 0.10 when no covariate is included. In contrast, none of “dist_AA”, “dist_CLAIRE_cor” and “dist_CLAIRE breadth” show significance either with or without covariates.

**Figure 4.**
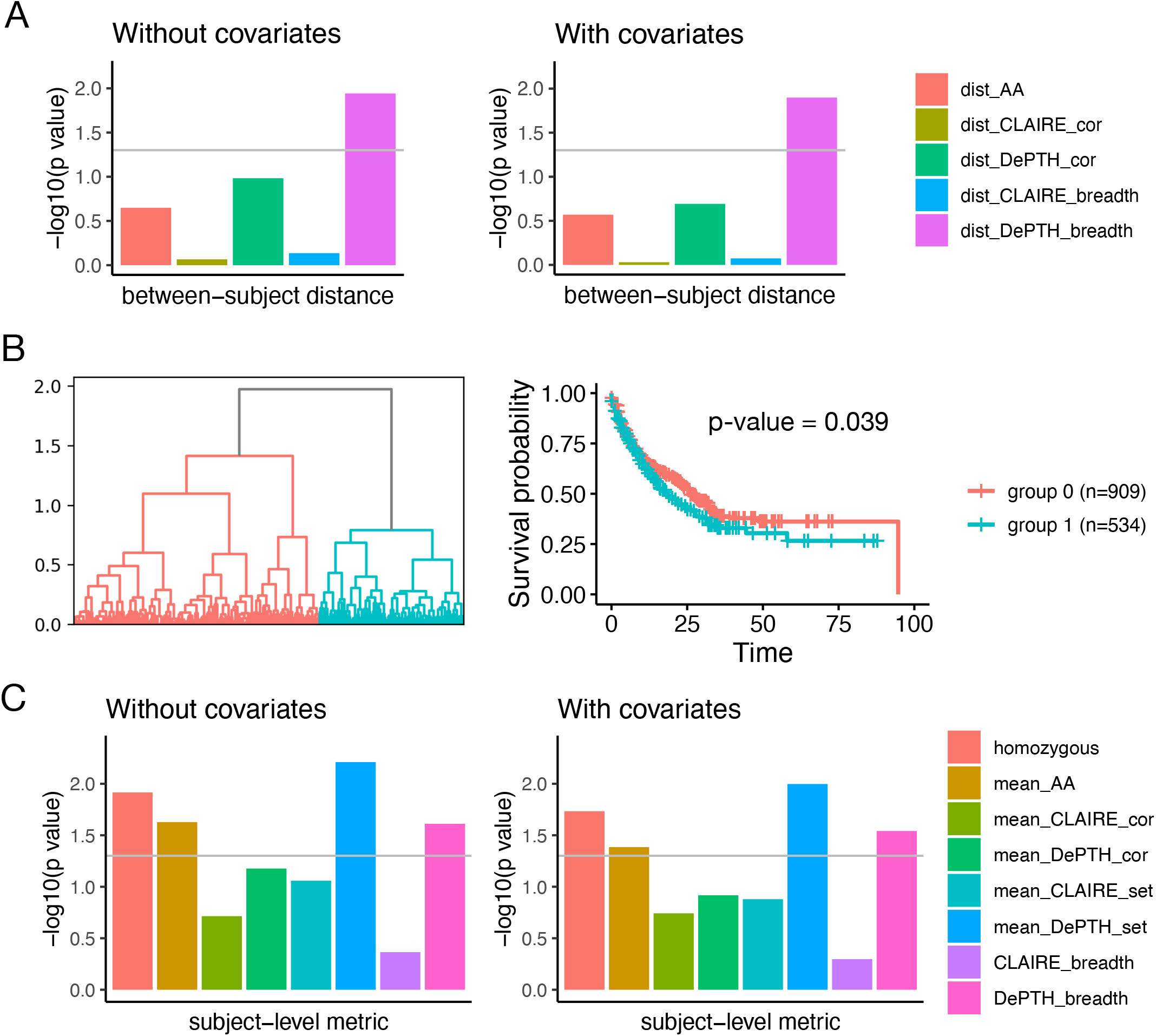
Association between HLA-based metrics and survival outcomes of 1,443 cancer patients [20]. (A) and (B) summarize the results using between-individual HLA-I distance metrics. (A)-log10(p value) of kernel regression using five different distance matrices with or without covariates. The covariates include age, log-transformed mutation burden, and drug type (CTLA-4, PD-1, or both). The grey horizontal line corresponds to p-value 0.05. (B) The dendrogram of hierarchical clustering where the distance is defined by “dist_DePTH_breadth”, and the survival curves of the two subgroups identified by cutting the dendrogram tree. The p value is computed from Cox proportional hazard regression. (C) -log10(p value) from Cox regression for the associations between individual-level HLA-I heterozygosity metrics and survival outcome, with or without the same set of covariates. The grey horizontal line corresponds to p-value 0.05.

The performance advantage over “dist_AA” indicates that our predicted scores may provide meaningful information in terms of association of HLAs with TCRs besides HLA pseudo sequences. The advantage over “dist_CLAIRE_cor” and “dist_CLAIRE_breadth” may be due to multiple factors. One is the flexibility of our model to make meaningful predictions for rare or previous unseen HLAs. Another factor is the limited diversity of HLA-I alleles among the McPAS data used for building CLAIRE model. The McPAS data only involve 35 different HLA-I alleles, which cover only 61 among all 146 different HLA-I alleles from all individuals after the additional HLA allele matching step. The third factor could be bias in the training data. The McPAS positive TCR-HLA pairs that are found through experiments might tend to have stronger binding, while the Emerson positive TCR-HLA pairs got from co-occurrence pattern may have weaker binding but reveal association in more general sense, regardless of the context of specific peptides.

We also conduct an un-supervised analysis to cluster patients by hierarchical clustering using the distance matrix formed by “dist_DePTH_breadth” (Figure 4(B)). The patients clearly separate into two groups whose survival time are significantlly different, with p value 0.039 from Cox proportional hazard regression.

#### Individual-level HLA-I heterozygosity metrics

Besides the between-individual distances, we also define eight individual-level HLA-I heterozygosity metrics to quantify the “variation” of HLA-I alleles within an individual, and assessed their associations with survival outcome through cox regression. “homozygous” is a simple metric, which has value 0 if an individual has two different alleles for all three HLA-I genes (HLA-A, HLA-B, and HLA-C), and 1 otherwise. For each individual, “DePTH_breadth” and “CLAIRE_breadth” are the total number of potentially cancer-related CD8+ TCRs that are predicted to be associated with at least one of her/his six HLA-I alleles based on the prediction scores by DePTH and CLAIRE, respectively (Figure 3(B)). For the other five metrics, we first calculate the distance between the two alleles of each HLA-I gene, and then take average across three HLA-I genes as the individual-level HLA-I heterozygosity. Among these metrics, “mean_DePTH_cor” and “mean_DePTH_set” are both based on the prediction scores from DePTH, where “mean_DePTH_cor” calculates the allele-level distance as 1 minus the Spearman’s correlation and “mean_DePTH_set” calculates set-level distance by Equation 1, except that the *A* and *B* sets are the sets of TCRs associated with each of the two HLA alleles. “mean CLAIRE_cor” and “mean CLAIRE set” are calculated similarly based on the prediction scores given by CLAIRE. “mean AA” uses the same allele-level distance as that for “dist_AA”. We run Cox regression on the 1,443 individuals to assess the association between each individual-level HLA-I heterozygosity metric and survival outcome. As shown in Figure 4(C), at p-value cutoff 0.05, when no covariate is included, “homozygous”, “mean AA”, “mean_DePTH_set”, and “DePTH_breadth” show significance and the significance remains when covariates (age, log-transformed mutation burden, and drug type) are included. These results suggest that DePTH is a useful alternative to quantify HLA similarities. Compared with “homozygous” which is 0/1, “mean_DePTH_set” and “DePTH_breadth” are continuous metrics and thus can provide a quantitative rather than categorical measurement.

## Discussion

We have proposed a neural network method DePTH to predict the associations between TCRs and HLAs based on their sequences. We demonstrate that two factors are important for model training: unbiased selection of training data from TCR and HLA co-occurrence patterns and representing an HLA allele by its sequence rather than treating it as a categorical variable.

We demonstrate an application of DePTH to define similarities of HLA-I alleles and show the resulting HLA metrics are associated with survival outcome of cancer patients undergoing cancer immunotherapy. A closely related question is whether the HLA metrics are associated with patient response to immunotherapy. The Chowell et al. [20] dataset do not have complete treatment response information and thus we turned to another dataset of cancer patients with melanoma and treated with PD-1 blockade, with or without prior-CTLA4 treatment [24]. In the PD-1 only group, the patients with low HLA heterozygosity metrics and mutation burdens tend to have worse outcome (Supplementary Figures 5&6), though it is hard to evaluate the association due to small sample size. Nevertheless, these results suggest a potential direction of using HLA-I and/or HLA-II heterozygosity metrics together with mutation burden as biomarkers for patient response to immunotherapy.

There are also many other situations where the results of DePTH can be very useful. For example, earlier works have shown that TCR data can be used to predict past infection (e.g., cytomegalovirus (CMV) [1] or SARS-COV-2 [3]). Since many TCRs are restricted to certain HLA alleles [1, 3, 7], incorporating HLA information could improve the prediction performance. This is a challenging task because HLAs are highly polymorphic and many HLA alleles are relatively rare in human population. Our DePTH method provides a solution to find HLA-specific TCRs and thus can help build HLA-specific TCR predictors for infection or other immune related conditions.

There are several directions that warrant further development. First, as more TCR data are accumulated, it is desirable to retrain DePTH with more data. Second, we have only considered TCR*β* chain in our method due to limitation of available data. As more paired TCR*α* and *β* chains are being generated (e.g., by single cell TCR data), it is possible to expand our model to include both TCR*α* and *β* chains.

Finally, furthering examination to the model and pinpointing the sequential structures that are important for the prediction might also provide new information for the biological mechanism under the binding of TCR and HLA sequences.

## Conclusions

We proposed a neural network method named DePTH to predict TCR-HLA associations based on their amino acid sequences. DePTH can make prediction for any TCR-HLA pairs provided sequence information is available. This property makes it possible to study rare HLAs, which is hard to do by examining co-occurrence patterns, due to the limited TCR repertoire data available.

To demonstrate the utilities of DePTH, we use the prediction scores of TCR-HLA associations given by DePTH to quantify functional similarities between HLA alleles, and further define between-individual HLA distances and individual-level heterozygosity metrics. We show that some of these distance metrics are significantly associated with survival outcomes of cancer patients.

## Methods

### Model training details

The number of negative pairs that we sample for each HLA allele is five times of the number of positive pairs. We randomly split both positive and negative pairs into three sets: 60% for training, 20% for validation and 20% for testing. The ratio of the number of positive pairs vs. that of negative pairs is 1:5 for all three sets.

In the neural network, the two dense layers, one for HLA and one for TCR, both have 64 nodes. The padded and numerically encoded CDR3 sequence is passed as input to the CNN layer in the format of a matrix, where each column is the encoding of one amino acid of the CDR3 sequence. In the CNN layer, we apply 8 filters, each with width 2 and height being the length of the encoded vector. These filters only slide in the direction across columns. After ReLU activation, one maxpooling layer with pooling size 2, stride 1 and no padding is applied. This CNN layer follows similar structure as the first of the two CNN layers in the model by [25].

The activation functions are ReLU (rectified linear unit), except for the output layer where sigmoid activation is used. The loss function is a binary cross-entropy loss, and it is optimized by the Adam optimizer with learning rate 0.0001. Since the number of positive versus negative TCR-HLA pairs has a ratio 1:5 in the training data, we assign a class weight 5:1 during training. We stop training if the AUC on validation data has not improved for 10 consecutive epochs, or if the number of training epochs reaches 300. We keep the model with the best validation performance before training stops and make prediction on test dataset.

### Hyperparameter choosing

We consider hyperparameters in two aspects, the encoding method for amino acids and the structure of the neural network after concatenating HLA and TCR parts together.

We consider four methods (one-hot, Blosum62, “Achtley” and “PCA”) to encode each amino acid to a numerical vector. One-hot encoding maps each amino acid into a vector of 0s and 1s, where the length of the vector equals the number of all possible amino acids (may extend to include wild card character “X”, gap character “.” or other characters when needed) and only the position corresponding to the given amino acid has value 1. Blosum62 [22] scores the similarity of two amino acids by the log odds of substitution between aligned sequences. “Achtley” refers to the five highly interpretable numeric patterns produced by Atchley et al. [26] through analysis on almost 500 amino acid indices representing different pysiochemical and biological properties. “PCA” refers to the top 15 principle components summarized by Beshnova et al. [25] from over 500 amino acid indices. See Section 3.2 of the Supplementary Materials for more details on encoding.

For neural network architecture after concatenating the outputs from HLA and TCR, we consider one or two layers feed forward neural network. If there is one layer, we consider three sizes: 16, 32, or 64. If there are two layers, we consider two options for their sizes: 32/16 or 64/16 for the first/second layer. A dropout layer can be added after the first dense layer. We consider three options, no dropout, dropout with probability 0.2 or dropout with probability 0.5. In total, we have 60 hyperparameter settings.

To choose a hyperparameter setting, we first run cross-validation on the set of TCR-HLA pairs formed by combining the training and validation data together. We randomly split this set of TCR-HLA pairs into two subsets with 3:1 ratio (i.e., 75% vs. 25%), while maintaining the 1:5 ratio of positive vs. negative pairs in each subset. Under each hyperparameter setting, we train the model on the 75% subset and stop training if either the AUC on the remaining 25% subset does not increase for 10 epochs or the number of epochs reaches 300, and record the highest AUC on the 25% subset. We repeat this procedure five times and record the average of the five recorded AUCs. The hyperparameter setting with the highest average AUC is chosen to train our model. The best hyperparameter settings chosen under different situations are listed in Supplementary Table 1.

### Compute between-individual distances

To compute each of between-individual distances “dist_AA”, “dist_DePTH_cor” and “dist_CLAIRE_cor”, we apply optimal transport to transform distances between HLA-I alleles to distances between individuals. More specifically, we represent one individual by 6 HLA-I alleles (possibly with duplicates), and then distance between two individuals can be characterized by a 6 by 6 matrix. Each entry of this matrix is the distance between two HLA alleles, and it can be considered as the cost to transfer between the two HLA alleles. The optimal transport method reduces this matrix to one number: the minimal cost to change all 6 HLA-I alleles from the first individual to the 6 HLA-I alleles of the second individual. The resulting minimal cost is taken as the between-individual distance.

The procedure of applying optimal transport is the same for “dist_AA”, “dist_DePTH_cor” and “dist_CLAIRE_cor”, except that for “dist_DePTH_cor” and “dist_CLAIRE_cor”, the distances between two HLA-I alleles are computed as 1 - Spearman’s correlation of TCR-HLA association scores given by DePTH and CLAIRE model, respectively. While on the other hand, for “dist_AA”, the distance between two HLA-I alleles *a*_1_ and *a*_2_ is calculated as

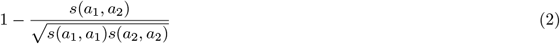

following the approach by Nielsen et al. [27], and here *s*(*a*_1_, *a*_2_) is the sum of the values on Blosum62 matrix corresponding to the amino acid from *a*_1_ and that from *a*_2_ on each aligned position of the pseudo sequences.

## Availability of data and materials

The TCR repertoire data of 666 individuals, which were generated by Emerson et al. [1], were downloaded from https://clients.adaptivebiotech.com/pub/Emerson-2017-NatGen in 2021, and the HLA data file for these individuals, which was generated by DeWitt et al. [7], was downloaded from Zenodo database (doi:10.5281/zenodo.1248193).

The file with survival outcome and HLA-I information for 1,535 advanced cancer patients treated with ICB, which was compiled by Chowell et al. [20], was downloaded from www.sciencemag.org/content/359/6375/582/suppl/DC1.

The whole exome sequencing data and the file with survival outcome for the 144 patients with melanoma and treated with PD1 ICB [24], were downloaded NCBI SRA, and the relevant files were located by dbGAP accession number phs000452.v3.p1.

The CLAIRE model, which was provided by Glazer et al. [11], was accessed from server at https://claire.math.biu.ac.il/Home.

Our data analysis pipeline is available at https://github.com/Sun-lab/DePTH_pipeline. Our software package is available at https://github.com/Sun-lab/DePTH [28] and also uploaded to the Python Package Index (PyPI).

Both repositories are licensed under the open source MIT License. A tutorial on package installation and usage is available at https://liusi2019.github.io/DePTH-tutorial/.

## Ethics approval and consent to participate

Not applicable.

## Consent for publication

All authors read and approved the final manuscript.

## Authors’ contributions

W.S. and S.L designed the study, with input from P.B. S. L. developed the software package and conducted the analysis. S.L., W.S. and P.B. wrote the paper.

## Additional Files

Additional file 1 — Supplementary Materials

Supplementary Methods and Results.

