## Supplementary Materials for "Neural Network Models for Sequence-Based TCR and HLA Association Prediction"

May 25, 2023

**8 Compute individual-level HLA-I and HLA-II metrics for patients in Liu 2019 study 9**

**9 Supplementary tables and figures 9**

### 1 Prepare (TCR, HLA) pairs from Emerson data

#### 1.1 Select public TCRs

The dataset that we use here contains information from 666 individuals and was initially reported by Emerson et al. [1] and later amended and re-analyzed by DeWitt et al. [2]. The TCR$\beta$ repertoire sequence data part of this dataset was generated by Emerson et al. [1] and downloaded in January 2021 from the link `https://clients.adaptivebiotech.com/pub/Emerson-2017-NatGen`. In our setting, a unique TCR$\beta$ chain is decided by two components, V gene and CDR3 amino acid sequence. We first go through the files of all individuals to record all the TCRs that appear in at least two individuals. We extract the V gene information on the V allele level and only keep the TCRs satisfying both the two criteria: (1) have frame type being "In"; (2) either have V allele information or V allele ties information. If there are V allele ties, we choose the first candidate allele listed in the ties.

Following one criterion in our earlier work [2], we remove those TCRs that are potential products of cross-contamination. After removal, we arrive at 8,739,207 unique public TCRs that each appears in at least two individuals.

#### 1.2 Threshold p-values for association

Based on the data generated by DeWitt et al. [2], there are 85 unique HLA-I alleles, 125 unique HLA-II alleles and 5 unique HLA-II haplotypes existing among the 666 individuals. We compute the association p-values between the 8,739,207 public TCRs and these 215 HLAs based on their co-occurrence pattern. The test used is one-sided Fisher's exact test. We choose a p-value cutoff corresponding to FDR (false discovery rate) 0.05. The FDRs at different p-value cutoffs are computed based on the corresponding average number of discoveries from 20 permutations. Viewing all pairs with p-value smaller than the p-value cutoff as positive pairs (with label 1), we get 20,582 associated (TCR, HLA) pairs, including 6,423 pairs involving HLA-I alleles and 11,037 pairs involving HLA-II alleles, excluding the pairs involving HLA-II haplotypes.

#### 1.3 Sample negative pairs

For building the model, we sample negative pairs for each unique HLA allele separately. For some HLA alleles, there are some individuals for whom no information on whether they have those HLA alleles or not exists. Given an HLA allele, we restrict on the individuals with known status on whether they have this HLA allele, and consider the set of TCRs that each appears in at least two of these individuals but does not form a positive pair with this HLA allele as the candidate pool for being selected. To match the prevalence among individuals with known existing or not status of this HLA allele, we first bin the TCRs involved in the positive pairs according to their prevalence and count the number of TCRs within each bin. The boundaries of these bins are formed by 10%, 20%, . . ., 90% quantiles of the prevalence of the TCRs involved in the positive pairs, with the maximal prevalence added as the rightmost boundary, and additional bins corresponding to exact prevalence 2, 3, . . ., 10 on the left. If 10 is smaller than the 10% quantile, the bins will correspond to prevalence 2, 3, . . ., 10, (10, 10% quantile], (10% quantile, 20% quantile], . . ., (90% quantile, maximum]. Otherwise, if 30% is the first quantile that is greater than 10, for example, then the bins will correspond to prevalence 2, 3, . . ., 10, (10, 30% quantile], (30% quantile, 40% quantile], . . ., (90% quantile, maximum]. For each bin, we count the number of TCRs forming positive pairs with this HLA allele and having prevalence falling into this bin, and sample five times many from the TCRs in the candidate pool that also have prevalence in this bin. All sampled TCRs for all bins are those forming the sampled negative (TCR, HLA) pairs. The reason of adding fine grid bins for the lower prevalence part is due to that majority of the TCRs in candidate pool have extremely low prevalence, such as 2 or 3, while those forming positive pairs with HLA are not, and we want to make the prevalence of the TCRs in sampled negative pairs as consistent as possible with that of those in positive pairs.

### 2 Features

#### 2.1 HLA pseudo sequence

We form the HLA pseudo sequences for 85 HLA-I alleles from Emerson data. For HLA-II alleles, we form the pseudo sequences for 250 HLA-II alleles, which include not only those directly from Emerson data, but also those with alpha and beta chains appearing in Emerson data. For example, HLA-DPAB*02:02_03:01 is an HLA-II allele formed by HLA-DPA1*02:02 and HLA-DPB1*03:01, and it does not exist in Emerson data, but both HLA-DPA1*02:02 and HLA-DPB1*03:01 exist as part of other HLA-II alleles in Emerson data. In this case, we form HLA pseudo sequence for HLA-DPAB*02:02_03:01 as well.

For each HLA class, the positions where HLA alleles are in contact with TCRs or antigens are used, which are 40 and 45 positions for HLA-I alleles and HLA-II alleles, respectively. To find these positions, we first include the 34 positions provided by Nielsen et al. [3] for HLA-I alleles and Karosiene et al. [4] for HLA-II alleles that are in contact with antigens and have amino acid polymorphism across different alleles, with the files of pseudo sequence for these positions provided by Reynisson et al. [5]. In addition, for HLA-I alleles, we find another 6 positions that each (1) has at least 10 records as a position where some HLA-I alleles contact the peptides or at least 10 records as a position where some HLA-I alleles contact the TCRs based on the information provided by DeWitt et al. [2], and (2) has amino acid polymorphism across the HLA-I alleles in Emerson data, where the amino acid sequences come from DeWitt et al. [2] and the IPD and IMGT/HLA database[6]. While for HLA-II alleles, for alpha chains, we add another another 7 positions with record number cutoff 3 and having polymorphism, and for beta chains, we add another 4 positions with record number cutoff 10 and having polymorphism.

#### 2.2 TCR sequence

Each TCR in our context contains two parts, the V gene and the CDR3. The sequence for each TCR is decided by the amino acid sequences in CDR1, CDR2 and CDR2.5 regions of the V gene and the amino acid sequence of CDR3. Schattgen et al. [7] provide a table for V genes and their corresponding CDR1, CDR2 and CDR2.5 sequences, but the original format of V genes from the resulting Emerson public TCRs of Supplementary Materials Section 1.1 is different from the format in this table. To solve this issue, first, we translate the V gene format from Emerson public TCRs to that in this table and use the translated format of V genes in input data files for DePTH. Second, when the input pairs are processed for DePTH, we get the corresponding CDR1, CDR2 and CDR2.5 amino acid sequences for each V gene from this table. Among the 65 unique V gene names that Emerson public TCRs involve, there are 5 V genes that each does not have a match in this table. For them, we translate them to a name "not_found" and then map it to sequences of character "." with the lengths of corresponding regions. When predictions are made on other data resources, the V genes that do not exist in this table are also mapped to the sequences of character "." in the same way.

### 3 Encoding

#### 3.1 CDR3 sequence padding

To pad all CDR3 amino acid sequences to the same length 27, which is the maximum CDR3 length from Emerson data, we follow the way used by Dash et al. [8], which starts padding right after position floor((L-5)/2) + 3 if the length L before padding is no greater than 10, and right after the 6th position if L is greater than 10.

##### 3.2 Encoding for characters that are not amino acids

Some characters in HLA pseudo sequence and CDR1, CDR2 and CDR2.5 amino acid sequences are not amino acid. Some HLA pseudo sequence contain the character "X". For encoding HLA pseudo sequence, when one hot encoding is used, the length of vector after encoding is set to be 21, with the extra position corresponding to "X". When Blosum62 encoding is used, the Blosum62 matrix used is the one with an additional row (and column) for "X". Under Atchley and PCA encoding, "X" is encoded as a vector of all 0s in both cases with corresponding vector length.

Some CDR2.5 and the padded CDR3 amino acid sequences contain ".". For encoding these two parts, when one hot encoding is used, the length of vector after encoding is extended to 21, with the extra position corresponding to ".". When Blosum62 encoding is used, the Blosum62 matrix is extended to have an additional column (and row) corresponding to "." with -4 as the first 20 values and 1 as the last value. Under Atchley and PCA encoding, "." is encoded as a vector of all 0s in both cases with corresponding vector length.

Some CDR1 and CDR2 amino acid sequences contain "." or "*". For encoding these two parts, we follow similar way as for CDR2.5 and CDR3. When one hot encoding is used, we extend the length of vector after encoding to 22. When Blosum62 encoding is used, both "." and "*" are encoded as a vector with -4 as the first 20 values and 1 as the last value. Under Atchley and PCA encoding, both "." and "*" are encoded as a vector of all 0s in both cases with corresponding vector length.

#### 4 Process solved TCR-pMHC-I complexes

All unique TCR-pMHC-I complexes with solved structures are extracted from Table 1 in the paper by Szeto et al. [9]. From these, we only keep those involving human HLA-I alleles from the 85 HLA-I alleles in Emerson data and exclude one complex with TCR$\beta$ chain V gene being TRGV8*01. For the complexes with CDR3 amino acid sequences not starting with "C" or not ending with "F", we add a letter "C" in front and a letter "F" at the end of the sequence. We get 54 pairs that are unique in terms of MHC, TRBV and CDR3$\beta$ columns. When getting prediction scores from DePTH trained on Emerson data, for TRBVs not found in the table for translating V genes to CDR1, CDR2 and CDR2.5, we rename them as "not_found" and later map them to sequences of character "." with the lengths of corresponding regions.

#### 5 Train DePTH on processed McPAS data

##### 5.1 Process McPAS datasets for training DePTH

The McPAS files for training, validation and test pairs used for CLAIRE model training are downloaded from the github repository provided by Glazer et al. [10]. We extract the "vb", "tcrb" and "mhc" information for all the pairs from these files. All pairs from training, validation and test combined involve 35 different HLA-I alleles. From these, 16 alleles belong to the 85 HLA-I alleles in Emerson data. From training pairs, we only keep those with HLA-I alleles belonging to one of these 16 alleles. The format of "vb" information from the downloaded McPAS files is different from the V gene names in the table by Schattgen et al. [7] for translating V genes to CDR1, CDR2 and CDR2.5. For V genes with "vb" information that can find a match in the table by Schattgen et al. [7], we convert their format, for example, by translating "TRBV01-01" to "TRBV1*01", "TRBV02-01" to "TRBV2*01" and "TRBV03-01" to "TRBV3-1*01". For V genes with "vb" that either do not have a match or appear as NA, we rename them as "not_found" and later map them to sequences of character "." with the lengths of amino acids sequences in corresponding regions. To keep it as consistent as possible with the data originally use for training CLAIRE, we keep all pairs after filtering by the 16 HLA-I alleles and do not remove duplicates in pairs. The same procedure is done for the validation and test pairs.

#### 5.2 Train DePTH on McPAS data

After processing the McPAS data, we arrive at 5090, 4691, 1102, 994, 1045 and 1041 pairs for positive training, negative training, positive validation, negative validation, positive test and negative test datasets, respectively. When training DePTH model on processed McPAS data, we assign class weight of positive v.s. negative to be 4691 v.s. 5090. When doing cross-validation to choose hyperparameter setting, we first combine the training and validation datasets together, and each time randomly split the combined set into two parts, one as new training set and one as new validation set, such that the new training set still has 5090 positive pairs and 4691 negative pairs. In other aspects, the training process is the same as that for training DePTH on Emerson data.

### 6 Get prediction scores for computing HLA-I diversity metrics for patients in Chowell 2018 study

#### 6.1 Obtain potentially cancer-related CD8+ TCRs

There are two data sources involved in obtaining the set of potentially cancer-related CD8+ TCRs. The first one contains the gene expression and TCR data for CD8+ T cells from eight cancer types provided by Zheng et al. [11] and the second one contains gene expression signatures related to neoantigen-reactivate or tumor-specific CD8+ T cells provided in multiple studies ([12], [13], [14], and [15]), which contains both up-regulated and down-regulated genes. The initial filtering of the CD8+ T cells is done by keeping the productive chains, removing the chains being "multi" and only keeping the beta chains with cdr3 length from 12 to 16 and cdr3 starting with C and ending with F. We only keep one beta chain per cell, and if one cell has multiple beta chains, the chain with the largest unique molecular identifier (UMI) counts is kept.

Treating each cancer type separately, for all the CD8+ T cells under a given cancer type, starting from the gene expression data, we first do PCA on the log-transformed normalized expression values from the expression count matrix involving the signature genes. Then, based on the first 20 principle components, we cluster the cells using Louvain method. We compute the gene set enrichment scores using GSVA for one set of down-regulated genes and three sets of up-regulated ones, and then scale the scores across T cells. For each T cell, we compute a score corresponding to the sets of up-regulated genes, by first computing the average of log-transformed normalized expression for each set, then scaling the results cross T cells, and finally taking an average of the scaled results and the scaled gene set enrichment scores for the sets of up-regulated genes. We compute a score corresponding to the sets of down-regulated genes in a similar way. For assigning T cell clusters to a label for being cancer-related (positive clusters) or not (negative clusters), for each cluster, we compute the median of the scores corresponding to the sets of up-regulated genes over the cells under this cluster and do the same for the scores corresponding to the set of down-regulated genes. The clusters with high final score for up-regulated genes and low final score for down-regulated genes are viewed as positive, and those with low final score for down-regulated genes and high final score for up-regulated genes are viewed as negative.

Across all the positive clusters under all cancer types, we extract the V gene and CDR3 amino acid sequence information from each CD8+ T cell and form a initial positive list of TCRs, where each TCR corresponds to one (V gene, CDR3) pair, and across all the negative clusters under all cancer types, we form another initial negative list following the same procedure. Multiple CD8+ T cells may share the same TCR, and some TCRs may appear multiple times in one list and some may appear in both lists. From the initial positive list, we only keep the TCRs with frequency in initial positive list greater than two times of the frequency in initial negative list. Further, to put a constraint on the influence of TCRs with extremely high frequency in the positive list, for TCRs each appearing more than 20 times, we only keep 20 copies of them. The resulting positive list of TCRs is our final list of potentially cancer-related CD8+ TCRs for building HLA-related metrics.

##### 6.2 Get scores from DePTH trained on Emerson data for (TCR, HLA) pairs between potentially cancer-related CD8+ TCRs and HLA-I alleles from patients in Chowell 2018 study

Among patients from Chowell 2018 study, some patients have HLA-I alleles that do not belong to the 85 HLA-I alleles from Emerson data. For each of these HLA-I alleles, if there are a group of HLA-I alleles from Emerson data that match it on the allele group level (i.e., they share the same two-digit HLA information), then we match it with the first allele in the group after ranking by allele name and use pseudo sequence of the first allele instead (e.g., HLA-A*02:02 does not exist in Emerson data, but HLA-A*02:01, HLA-A*02:05 and HLA-A*02:06 do. In this case, we match HLA-A*02:02 with HLA-A*02:01 and take the pseudo sequence of HLA-A*02:01 as that for HLA-A*02:02). We exclude patients with HLA-I alleles that do not belong to the 85 alleles from Emerson data and we cannot find a match for, and this reduces the total sample size from 1535 to 1443, with the 1443 individuals involving 146 different HLA-I alleles.

The format of V genes from potentially cancer-related TCRs has difference from those used by DePTH model, and this difference is resolved by adding "*01" to the end of each V gene, for example, changing "TRBV2" to "TRBV2*01" and "TRBV3-1" to "TRBV3-1*01". The pairs formed by HLA-I alleles from the patients and potentially cancer-related TCRs are submitted to DePTH model trained on Emerson data to get prediction scores. For each HLA-I allele outside the 85 alleles from Emerson data but we can find a match for, the pseudo sequence of the matched HLA-I allele is used instead.

##### 6.3 Get scores from CLAIRE for (TCR, HLA) pairs between potentially cancer-related CD8+ TCRs and HLA-I alleles from patients in Chowell 2018 study

The V genes from the potential cancer-related TCRs are transformed to the format as in the CLAIRE McPAS-based training, validation and test data sets provided in the CLAIRE github repository. For example, TRBV3-1 is converted to TRBV03-01 and TRBV10-1 is converted to TRBV10-01. For the V genes that only come with family level gene name, for instance, TRBV13, we convert it by adding "-01" at the end and match it to TRBV13-01.

There are 146 unique HLA-I alleles from the 1,443 cancer patients kept from the Chowell 2018 study. Due to the limited coverage of McPAS data used by CLAIRE in terms of unique HLA-I alleles (there are only 35 unique HLA-I alleles that each appears in at least one of McPAS training, validation, and test sets), only 22 out of these 146 HLA-I alleles appear in McPAS data. There are 39 out of the other HLA-I alleles that each has a corresponding group of HLA alleles in McPAS data at two-digit level. For these 39 HLA-I alleles, we matched them to the first allele in the corresponding group. For example, HLA-A01:02 does not appear in the McPAS data, but HLA-A01:01 does, and in this case, we replace HLA-A01:02 with HLA-A01:01. We form pairs between all potentially cancer-related TCRs and all 146 HLA-I alleles (with the 39 HLA-I alleles replaced by their matched alleles in McPAS data) and upload them to the CLAIRE model held on server `https://claire.math.biu.ac.il/Home` to get prediction scores. The input format for CLAIRE allows components from TCR$\alpha$ chain, J gene for TCR$\beta$ chain and T cell types, while our pairs do not have information for these fields. We convert the pairs according to the format in the example files on CLAIRE server by filling "UNK" for tcra and leaving NA for va, ja, jb and T cell type columns.

#### 7 Get prediction scores for computing HLA diversity metrics for patients in Liu 2019 study

##### 7.1 Obtain potentially cancer-related CD8+ TCRs and CD4+ TCRs

The set of potentially cancer-related CD8+ TCRs for computing HLA-I diversity metrics for patients in Liu 2019 study is the same as that used for patients in Chowell 2018 study, and the procedure of obtaining these CD8+ TCRs is in Supplementary Materials Section 6.1. The procedure of obtaining potentially cancer-related CD4+ TCRs is similar to that used for obtaining CD8+ TCRs as listed in Supplementary Materials Section 6.1, with difference in the choice of signature genes.

##### 7.2 Get scores from DePTH trained on Emerson data for (TCR, HLA) pairs between potentially cancer-related CD8+ TCRs and HLA-I alleles from patients in Liu 2019 study

The data from Liu 2019 study contain whole-exome sequencing (WES) results on matched pretreatment tumor sample and normal tissue for 144 patients diagnosed with advanced melanoma and treated with anti-PD1 ICB. We used OptiType [16] for HLA-I typing. When deciding HLA-I alleles, the OptiType result from tumor sample is chosen for each patient. Some patients have HLA-I alleles that do not belong to the 85 HLA-I alleles from Emerson data, and we try finding match for these HLA-I alleles in the same approach as listed in Supplementary Materials Section 6.2. We exclude patients with HLA-I alleles that do not belong to the 85 alleles from Emerson data and we cannot find a match for, and this reduces the total sample size from 144 to 143. The procedures of reformatting TCRs and submitting to DePTH model to get scores are also the same as those listed in Supplementary Materials Section 6.2.

##### 7.3 Get scores from DePTH trained on Emerson data for (TCR, HLA) pairs between potentially cancer-related CD4+ TCRs and HLA-II alleles from patients in Liu 2019 study

We used HLA-HD [17] for HLA-II typing, with parameter -m being 60 and -c being 1 in the command line. Some patients have HLA-II alleles that do not belong to those from Emerson data. For each of these HLA-II alleles, if there are a group of HLA-II alleles from Emerson data and matching it on the allele group level (i.e., they share the same two-digit HLA information), then we match it with the first allele in the group after ranking by allele name and use pseudo sequence of the first allele instead (e.g., HLA-DQA1*01:10 does not exist in Emerson data, but HLA-DQA1*01:01, HLA-DQA1*01:02, HLA-DQA1*01:03, HLA-DQA1*01:04 and HLA-DQA1*01:05 do. In this case, we match HLA-DQA1*01:10 with HLA-DQA1*01:01 and take the pseudo sequence of HLA-DQA1*01:01 as that for HLA-DQA1*01:10). HLA-HD results have difference between tumor sample and normal tissue on 9 patients. For each of these 9 patients, when deciding the results from which sample to use, we first look at whether the HLA-II alleles all appear in Emerson data or have a match from Emerson data. If both tumor and normal sample satisfy this criterion, the results from tumor sample is chosen. Otherwise, if only the normal sample satisfy this criterion, the normal sample is chosen. In addition, for one patient, the HLA-HD typing on tumor sample was not finished within 30 days, and the results from normal sample are chosen. We exclude patients with HLA-II alleles that do not belong to the HLA-II alleles from Emerson data and we cannot find a match for, and this reduces the total sample size from 144 to 120.

HLA-II class contains three major types of proteins, HLA-DR, HLA-DP and HLA-DQ, and each of them contains an $\alpha$-chain and a $\beta$-chain. The HLA-II proteins from Emerson data come in a format of combined $\alpha$- and $\beta$-chains. To get scores from DePTH model trained on Emerson data, we convert the format of HLA-IIs from Liu 2019 data to that as in Emerson data. So, for each patient, we list all combinations of DPA1s and DPB1s as well as all combination of DQAs and DQBs. For example, if a patient has HLA-HD output being DRB1*03:01, DRB1*11:01, DQA1*05:01, DQA1*05:05, DQB1*02:01, DQB1*03:01, DPA1*01:03, DPA1*02:01, DPB1*01:01 and DPB1*02:01, then for this patient, there are two HLA-DR proteins which are HLA-DRB1*03:01 and HLA-DRB1*11:01, four HLA-DP proteins which are HLA-DQAB*05:01_02:01, HLA-DQAB*05:01_03:01, HLA-DQAB*05:05_02:01 and HLA-DQAB*05:05_03:01, and four the HLA-DQ proteins which are HLA-DPAB*01:03_01:01, HLA-DPAB*01:03_02:01, HLA-DPAB*02:01_01:01, HLA-DPAB*02:01_02:01. The reason that HLA-DR proteins only show up in the format of $\beta$-chain is due to that the $\alpha$-chain does not have polymorphisms in the peptide binding part.

### 8 Compute individual-level HLA-I and HLA-II metrics for patients in Liu 2019 study

The procedure of individual-level HLA-I metrics is the same as that used for patients in Chowell 2018 study. For individual-level HLA-II metrics, we follow similar procedure as that used for HLA-I metrics, with the difference in the treatment for HLA-DP and HLA-DQ. When computing mean_AA, for HLA-DP, since it has four alleles, we first calculate the distance between any pairs of two alleles out of the four, and then take the average of the distances from the six alleles as the distance value from HLA-DP. We do the same for HLA-DQ. For HLA-DR, since it only has two alleles, we compute the distance between the two alleles. Finally, we take the average of the three distance values from HLA-DR, HLA-DP and HLA-DQ as the individual-level HLA-II metric. The procedure of computing mean_DePTH_cor and mean_DePTH_set is similar to that for computing mean_AA. DePTH_breadth is the total number of cancer-related CD4+ TCRs that are predicted to be associated with at least one of the 10 HLA-II alleles from the patient.

### 9 Supplementary tables and figures

Table S1: Hyperparameter settings chosen from cross-validation for DePTH under different data sets. The first column shows the data that the model is trained on, and the second to last columns each gives the encoding method, the number of dense layers after concatenation and before final output layer, the sizes of dense layers, and the dropout probability. The first two rows correspond to training on Emerson pairs involving HLA-I and those involving HLA-II, separately. The third to the fifth rows each corresponds to the leave-one-out experiments based on Emerson pairs when HLA-B∗08:01, HLA-B∗07:02, and HLA-C∗07:01 is left in the test pairs, respectively. The last row corresponds to the case when DePTH is trained on the subset of McPAS pairs used for training CLAIRE that involve HLA-I alleles from the 85 HLA-I alleles in Emerson data.

| data | encoding | n dense layers | size of dense layer | dropout |
|---|---|---|---|---|
| Emerson HLA-I | one hot | 2 | [64, 16] | 0.2 |
| Emerson HLA-II | one hot | 2 | [64, 16] | 0.2 |
| Emerson HLA-B∗08:01 | pca | 1 | [64] | 0.5 |
| Emerson HLA-B∗07:02 | pca | 1 | [64] | 0.5 |
| Emerson HLA-C∗07:01 | pca | 2 | [64, 16] | 0.2 |
| McPAS HLA-I | pca | 2 | [64, 16] | 0.2 |

Table S2: Performance of DePTH trained on Emerson data and tested on Emerson test pairs, for HLA-I and HLA-II classes separately.

| HLA class | AUC | recall | specificity |
|---|---|---|---|
| HLA-I | 0.82 | 0.63 | 0.86 |
| HLA-II | 0.79 | 0.63 | 0.80 |

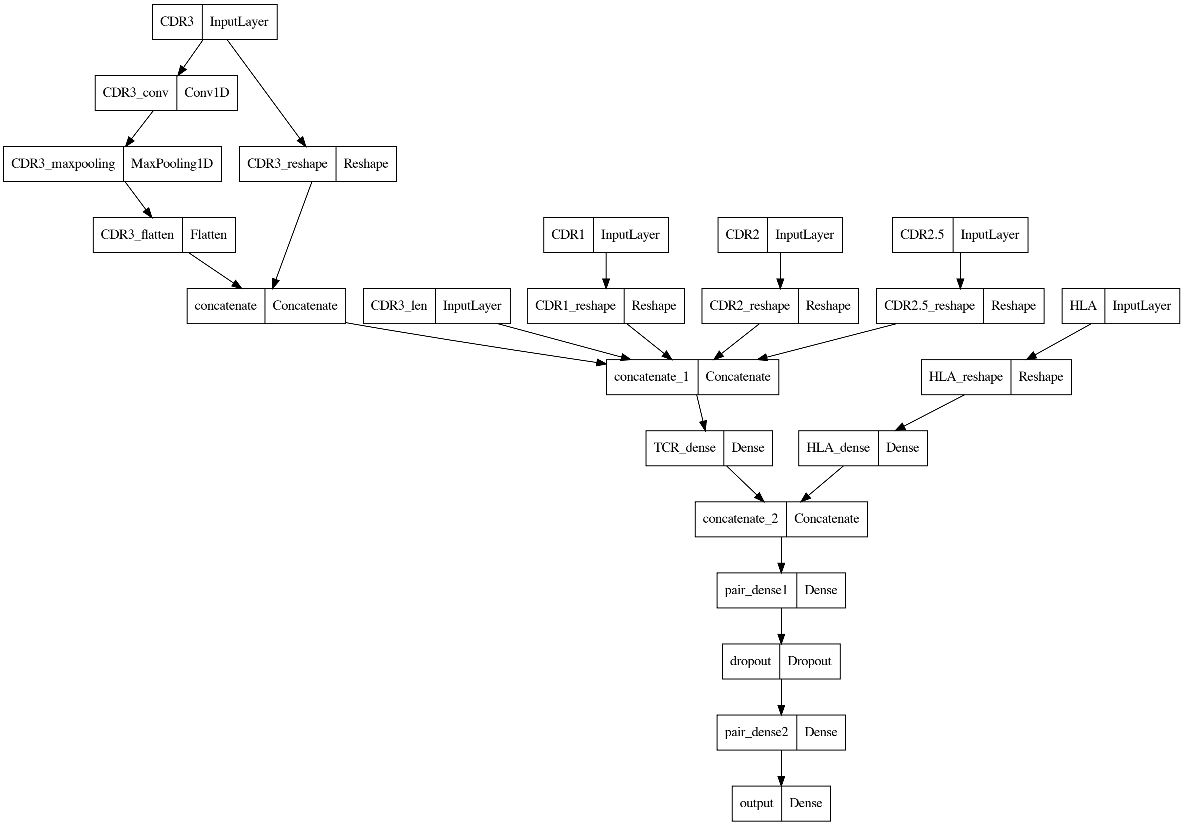

Figure S1: DOT graph showing the architecture of DePTH model in the situation of Emerson HLA-I or HLA-II data. The hyperparameters were chosen from cross-validation. A more detailed version of the graph with the shape of each layer can be found at `https://github.com/Sun-lab/DePTH_pipeline/blob/main/figures/depth_draft/supp6_hla_i_model_plot_w_size.pdf` for HLA-I and `https://github.com/Sun-lab/DePTH_pipeline/blob/main/figures/depth_draft/supp6_hla_ii_model_plot_w_size.pdf` for HLA-II.

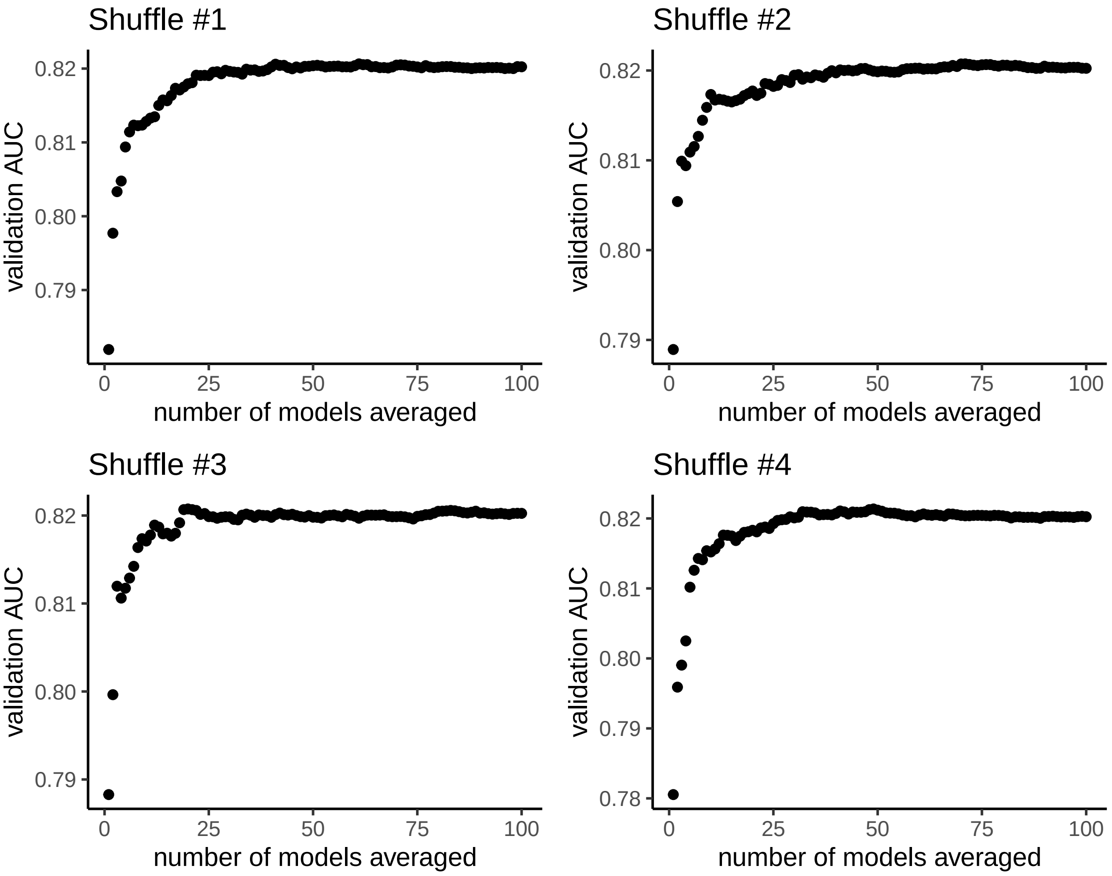

Figure S2: Validation AUC based on prediction scores averaged from the first $n$ models v.s. $n$, under four shuffles of 100 models trained using 100 different sets of random seeds. The models are trained on (TCR, HLA) pairs from Emerson data that involve HLA-I alleles, with the hyperparameter setting chosen from cross-validation, as listed in the first row of Supplementary Table 1.

Table S3: Performance of DePTH on leave-one-out experiments

| HLA allele | AUC | recall | specificity |
|---|---|---|---|
| HLA-B*08:01 | 0.64 | 0.26 | 0.92 |
| HLA-B*07:02 | 0.69 | 0.22 | 0.95 |
| HLA-C*07:01 | 0.69 | 0.31 | 0.92 |

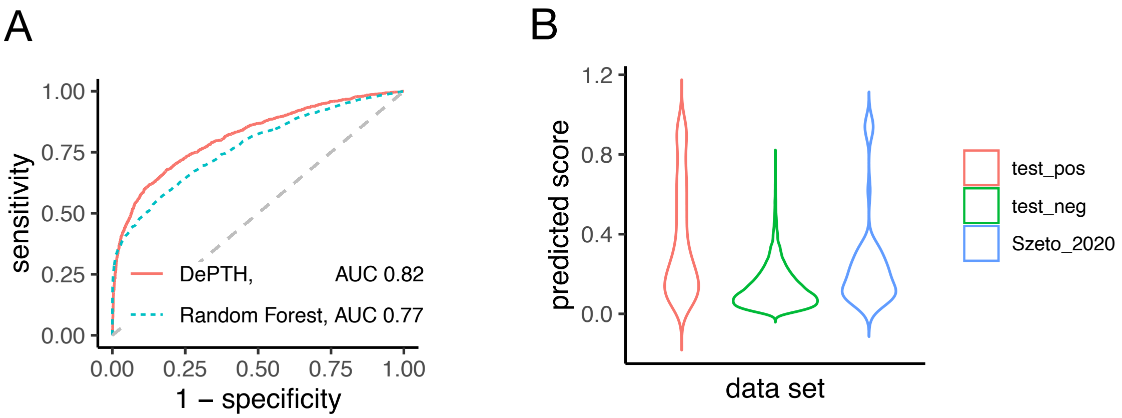

Figure S3: (A) ROC curves of DePTH prediction and random forest prediction on test data (part of Emerson data) for HLA-I. (B) Violin plots for the TCR-HLA association scores predicted by random forest on three sets of TCR-HLA pairs: the positive pairs of test data, the negative pairs of test data, and the 54 external positive TCR-HLA pairs from solved TCR-pMHC-I structures as listed in [9]. The scores are given by random forest trained on encoded Emerson training data for HLA-I and the encoding method is one-hot-encoding on the amino acid sequences.

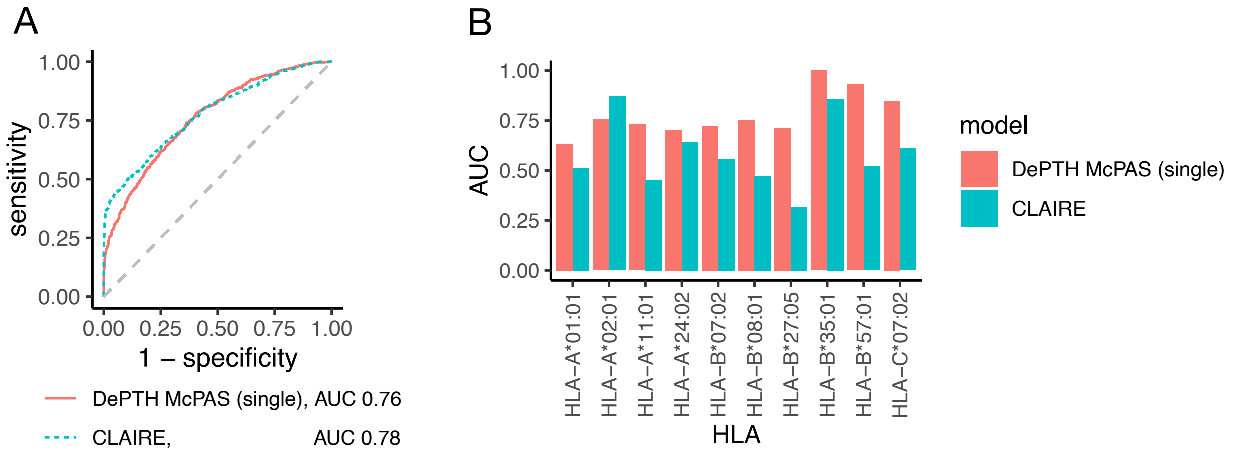

Figure S4: Comparison on the prediction performance on McPAS test data from one single DePTH model trained on McPAS data and that from CLAIRE model, with all training, validation and test pairs restricted to those involving HLA-I alleles from Emerson data. (A) ROC curves of predicton scores on McPAS data. (B) Comparison of allele-wise AUCs in McPAS test data between one single DePTH model trained on McPAS data and CLAIRE model.

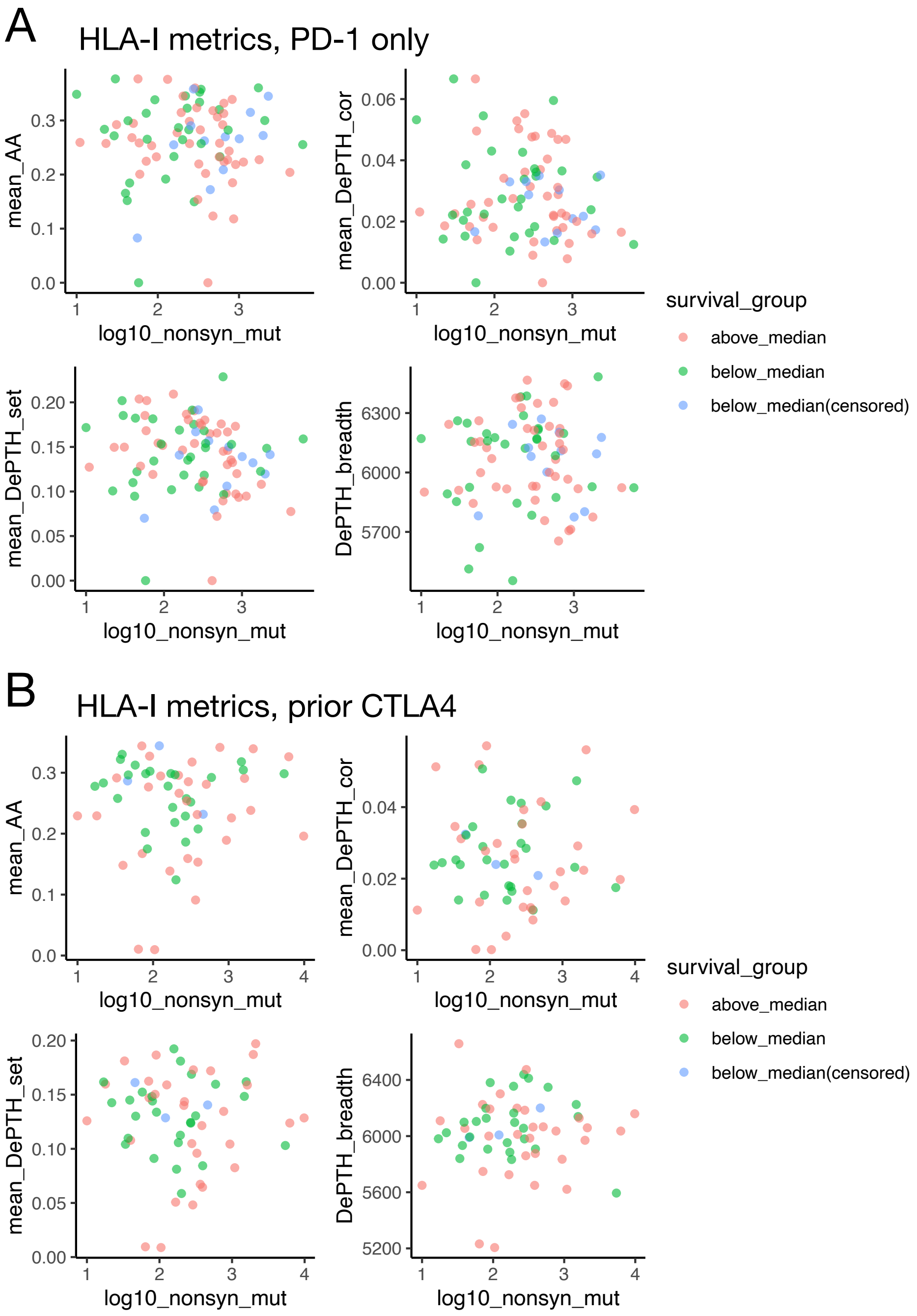

Figure S5: Scatterplots of HLA-I metrics v.s. log10(nonsynonymous mutations), colored by survival outcome, based on a cohort of patients with melanoma treated with anti-PD1 ICB [18] after processing. In each subset, the cutoff is chosen as the median observed survival time (can be from event or censoring), and then the patients are separated into three groups, the group with observed survival time above median (above_median), the group with survival time from event below median (below_median) and the group with survival time from censoring below median (below_median(censored). (A) The subset of patients who received no CTLA4 treatment prior to receiving PD-1 treatment (n=84). (B) The subset of patients who received prior CTLA4 treatment (n=59).

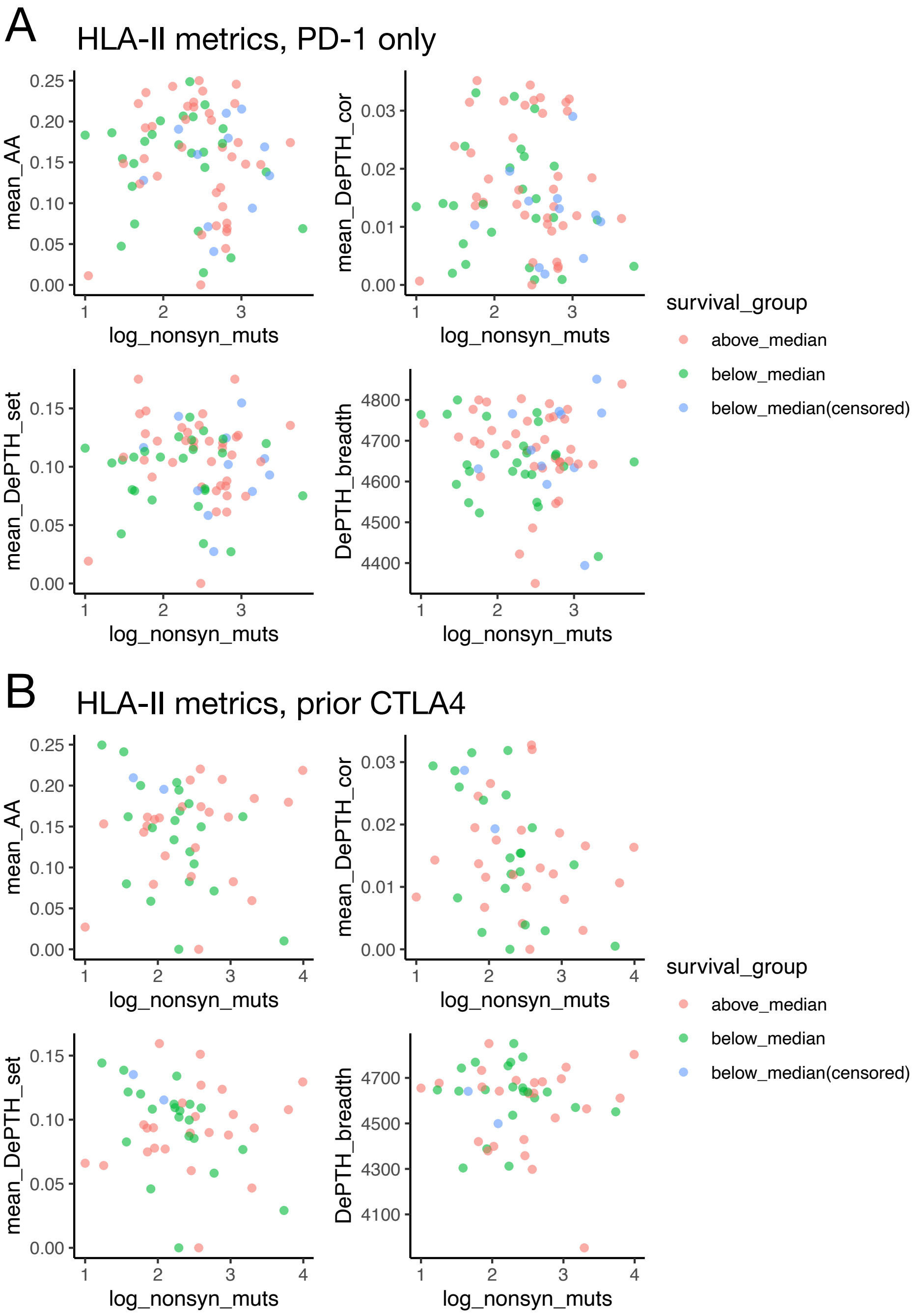

Figure S6: Scatterplots of HLA-II metrics v.s. log10(nonsynonymous mutations), colored by survival outcome, based on a cohort of patients with melanoma treated with anti-PD1 ICB [18] after processing. In each subset, the cutoff is chosen as the median observed survival time (can be from event or censoring), and then the patients are separated into three groups, the group with observed survival time above median (above_median), the group with survival time from event below median (below_median) and the group with survival time from censoring below median (below_median(censored). (A) The subset of patients who received no CTLA4 treatment prior to receiving PD-1 treatment (n=73). (B) The subset of patients who received prior CTLA4 treatment (n=47).
